# Characterization of Reactive Oxygen Species Signaling changes in a cell culture model of skeletal muscle ageing, and its application to screening pharmacokinetically-relevant exposures of dietary polyphenols for bioactivity

**DOI:** 10.1101/2021.11.20.469396

**Authors:** N. Hayes, M. Fogarty, L. Sadofsky, H.S. Jones

**Affiliations:** Department of Biological and Marine Sciences, University of Hull, Hull, HU6 7RX; Leeds Trinity University, Leeds, LS18 5HD; Centre for Atherothrombotic and Metabolic Research, Hull York Medical School, Hull, HU6 7RX; Institute of Cancer Therapeutics, University of Bradford, Bradford, BD7 1DP

## Abstract

Age-related frailty is a significant health and social care burden, however treatment options are limited. There is currently a lack of suitable cell culture model for screening large numbers of test compounds to identifying those which can potentially promote healthy skeletal muscle function. This paper describes the characterization of reactive oxygen and nitrogen species (RONS) signalling changes in young and aged myoblasts and myotubes using the C_2_C_12_ cell line, and the application of aged myoblast and myotube cultures to assess the effect of dietary polyphenols on RONS signalling. Aged myoblasts and myotubes were observed to have significantly increased reactive oxygen species levels (p<0.01 and p<0.001 respectively), increases in nitric oxide levels (p<0.05 for myoblasts and myotubes), and lipid peroxidation markers (p<0.05 for myoblasts and myotubes). A panel of nine polyphenols were assessed in aged myoblasts and myotubes using concentrations and incubation times consistent with known pharmacokinetic parameters for these compounds. Of these, although several polyphenols were seen to reduce single markers of RONS signalling, only kaempferol and resveratrol consistently reduced multiple markers of RONS signalling with statistical significance in both cell models. Modulation of cellular enzymatic antioxidant activities (superoxide and catalase) was assessed as a possible mechanism of action for these polyphenols, and although both superoxide and catalase activities were significantly reduced in aged (versus young) myotubes (p<0.01 and p<0.05 respectively), no effect of polyphenol treatment on these enzyme activities were observed. Overall, this research has shown the utility of the C_2_C_12_ model, as both myoblasts and myotubes, as a suitable cell model for screening compounds for modulating RONS signalling in aged muscle, and that resveratrol and kaempferol (using pharmacokinetically-informed exposures) can modulate RONS signalling in skeletal muscle cells after an acute exposure.

## Introduction

Age-related muscle dysfunction is a major healthcare issue in elderly patient, and is associated with significant reductions in functional capacity, independence, and quality of life (Int. J. Endocrinol., 2021, 5563960). It is also associated with increased risk of chronic debilitating disorders (such as obesity), falls, hospitalization and mortality (Int. J. Endocrinol., 2021, 5563960). Currently, the management of age-related muscle dysfunction consists of diet and lifestyle changes (e.g., increased physical exercise, intake of protein, vitamin D and calcium, weight management), and symptomatic treatments such as pain relief (Ther. Adv. Muscluloskelet Dis., 2021, 13, 1759720×211009018). These approaches have had some success in improving patient quality of life, however this outcome is not consistent for all patients (Ther. Adv. Muscluloskelet Dis., 2021, 13, 1759720×211009018). Due to the increasing life expectancy, the number of elderly patients suffering with age-related loss of muscle function is predicted to also increase (Int. J. Endocrinol., 2021, 5563960). Thus, identifying new treatments for age-related muscle dysfunction is an important clinical need.

Reactive oxygen and nitrogen species (RONS) signaling is well-accepted to have important physiological signaling functions in healthy skeletal muscle, and its dysregulation is also strongly implicated in age-related skeletal muscle dysfunction (J. Physiol., 2016, 594, 5081; Biogerontology, 2020, 21, 475). This makes measurements of RONS signaling important biomarkers, and therapeutic targets of skeletal muscle dysfunction. The current state of play in this area has recently been reviewed by Thoma et al (Biogerontology, 2020, 21, 475), highlighting current attempts to modulate RONS signaling have produced mixed results, dependent on the model systems and dosing regimes used (Biogerontology, 2020, 21, 475).

The dietary polyphenol resveratrol has shown encouraging results in terms of reducing RONS signaling dysfunction and improvement in aged muscle function in both cell culture and mouse models (Sci. Rep., 2015, 28, 8093; J. Nutr. Biochem., 2017, 50, 103; J. Physiol. Sci., 2018, 68, 681). There are some limitations to these studies, particularly the use of supra-physiological concentrations of polyphenols for supra-physiological exposure times, this lack of reflection of the pharmacokinetics of polyphenols is becoming a more recognized issue within dietary polyphenol research (J. Agric. Food Chem., 66, *7857;* J. Agric. Food Chem., 66, 8221). Taken together, this suggests that dietary polyphenols such as resveratrol could have therapeutic application to age-related muscle dysfunction, however a more thorough and physiologically-relevant assessment of these compounds needs to be done.

This study, using the C_2_C_12_ murine myotube model described by Sharples et al (J. Cell. Physiol., 2010, 225, 240), characterizes changes in RONS signaling between young and aged myoblasts and myotubes, which has not been previously reported for this model system. A range of dietary polyphenols, using pharmacokinetics-informed concentrations and exposure durations, were assessed for effects on RONS signaling in aged cells, to identify those which ameliorate age-related changes in these markers. Those polyphenols which alter ROS signaling in the screening assays were then further characterized for effects on ROS signaling in aged myoblasts and myotubes.

## Materials and Methods

### Materials

All materials were purchased from Sigma-Aldrich (Poole, UK) unless otherwise stated.

### Cell culture

Murine myoblasts (C_2_C_12_) were purchased from Public Health England at Passage 13 and seeded into either T-75 flasks (Biolite, Thermofisher Scientific), 6 well plates (Biolite, Thermofisher) or 96 well plates (Biolite, Thermofisher Scientific) at a density of 2.75×10^4^ cells/cm^2^. Myoblasts were cultured in high glucose Dulbecco’s Modified Eagle Medium (DMEM) containing L-glutamine, 10% v/v FBS, 100 U penicillin, 100 µg/mL streptomycin, and 12.5µg/mL amphotericin B, to a confluency of 80%, and reseeded using a 0.5% trypsin solution. To differentiate myoblasts into myotubes, cells were serum-starved (DMEM media containing L-Glutamine, 2% v/v horse serum, 100 U penicillin, 100 µg/mL streptomycin, and 12.5µg/mL amphotericin B) for 5-7 days, with media refreshed every 2 days.

To age myoblast and myotube cultures, myoblasts were continuously cultured for 50 population doublings as outlined by Sharples et al 2010. Myoblasts less than passage 20 were defined as “young”, with those above passage 30 as “aged”. For aged myotubes, myoblasts were cultured to the appropriate passage number, and were then differentiated as described above.

### Assessment of myotube length and width

C_2_C_12_ myotubes were cultured as described above and imaged by Olympus IX71 Inverted Fluorescence Microscope (10x UPlanXApo Objective), and the thickness of myotubes was determined using the ImageJ software (Version 1.53m). Five 6-well plates were used for each sample group 3 images per well, 5 fibres per image, a mean of three measurements of each myotube diameter was taken and one centermost measurement was taken for length.

### FOX assay

Lysates of myoblasts and myotubes grown in T-75 flasks were prepared after washing cultures with phosphate buffered saline (PBS). Lysates were stored at -20°C until analysed. Cell lysate or hydrogen peroxide standards (80 µL, 0-5 µM) were added into wells of a 96-well plate, with FOX reagent (7.6 mg of xylenol orange, 9.8 mg of ammonium ferrous sulphate and 1.8217 g of sorbitol together in 100 mL of 25 mM sulphuric acid, 120 µL per well) added to each well, and incubated for 30 min in the dark. Absorbance was measured at λ=560 nm using a Tecan infinite 200 mx-pro plate reader. Lipohydroperoxide concentrations were calculated for each sample using the hydrogen peroxide standard curve, and then normalized to the protein content of each lysate using Bradford assay.

### TBARS assay

Lysates of myoblasts and myotubes grown in T-75 flasks were prepared after washing cultures with PBS. Lysate was collected using a cell scraper and suspended in 1.5 mL PBS and stored at -20°C until analysed. The following steps were performed on ice and light shielded wherever possible.1 mL of cell lysate suspension was added to 100 µL of 6% Perchloric acid (PCA) and incubated for 10 mins, Supernatant was spun off on max speed, 4°C for 5 mins. Supernatant was then neutralised with 10 µL of 6 M Potassium Hydroxide Supernatant was spun off on max speed, 4°C for 5 mins and filtered through a 0.22 µm filter. 40 µL of sample or 1,1,3,3-Tetraethoxypropane (TEP)/ 40% ethanol standard (0-10 µM) were combined with 200 µL 438 mM Phosphoric acid, 200 µL 41.6 mM Thiobarituric acid (TBA) and 350 µL 18.2 MΩ Water. Samples and standards were then heated at 90°C for 60 min in the dark. Samples were then transferred to amber HPLC vials and sealed with a crimp top lid.

Chromatographic separation was achieved using an Agilent G1311A quaternary pump, Agilent G1367E1260 Infinity High Performance Autosampler and Agilent 1100 Series G1316A COLCOM Column Compartment connected to an Agilent 1260 Infinity Fluorescence Detector. An Agilent ZORBAX Eclipse Plus C18 column (5 µm pore size, 4.6 × 250 mm, Agilent Technologies, Cheshire, UK) was used for separating the analytes as detailed below: Solvent A consisted of Phosphate buffer (70% 25 mM KH_2_PO_4_ : 30% 25 mM Na_2_HPO_4_) and Solvent B was Acetonitrile. The Column was maintained at 25°C, Flow rate was 1 mL/min, injection volume was 25 µL, starting conditions were equilibrated over 10 min. The method began at 25% Solvent B from 0-1.5 min, rising to 50% over 1.51-3 min, holding at 50% until 4 min then dropping at 4.01 min, finally holding at 25% until 6 min. Using FLD λex = 532 nm and λem = 553 nm, TBARS were detected at RT = 3.5 min.

### DHE assay

Myoblasts and myotubes were cultured in a 96-well plate as described above, leaving the edge wells of the plate free of cells. Once cells had reached the desired confluence or differentiated into myotubes, cell culture media was removed from the wells and replaced with PBS, containing 10 µM Dihydroethidium (DHE, Abcam, UK, 100 µL per well). The edge wells were filled with 100 µL PBS. The well plates also contained cell only control wells (no DHE added), DHE only control wells (no cells), and positive control wells (10 µM DHE, 1 mU xanthine oxidase, 1 mM xanthine, in PBS). The plate was incubated at 37°C in a BMG Labtech fluorostar omega plate reader, with fluorescence intensity of each well measured each minute (using orbital averaging) for 30 min at λ_ex_ 544 nm and λ_em_ 590 nm. At the end of the assay, the wells were emptied and replaced with 100 µL of PBS, and frozen before quantification of protein content for each well using the Bradford assay. The data was analysed by calculating the linear rate of DHE fluorescence increase for each well, which was then corrected for protein content and the rate of the positive control samples. This was done to ensure reproducibility between different plates and experimental days.

### DCF assay

Myoblasts and myotubes were cultured in a 96-well plate as described above, leaving the edge wells of the plate free of cells. Once cells had reached the desired confluence or differentiated into myotubes, cell culture media was removed from the wells and replaced with PBS, containing 10 µM Dichlorofluorescein (DCF, Abcam, UK, 100 µL per well). The edge wells were filled with 100 µL PBS. The well plates also contained cell only control wells (no DHE added), DHE only control wells (no cells), and positive control wells (10 µM DCF, 1 mU xanthine oxidase, 1 mM xanthine, in PBS). The plate was incubated at 37°C in a BMG Labtech fluorostar omega plate reader, with fluorescence intensity of each well measured each minute (using orbital averaging) for 30 min at λ_ex_ 485 nm and λ_em_ 520 nm. At the end of the assay, the wells were emptied and replaced with 100 µL of PBS, and frozen before quantification of protein content for each well using the Bradford assay. The data was analysed by calculating the linear rate of DCF fluorescence increase for each well, which was then corrected for protein content and the rate of the positive control samples. This was done to ensure reproducibility between different plates and experimental days.

### DAF assay

Myoblasts and myotubes were cultured in a 96-well plate as described above, leaving the edge wells of the plate free of cells. Once cells had reached the desired confluence or differentiated into myotubes, cell culture media was removed from the wells and replaced with PBS, containing 2 µM 4,5-Diaminofluorescein diacetate (DAF, Abcam, UK, 100 µL per well). The edge wells were filled with 100 µL PBS. The well plates also contained cell only control wells (no DAF added), DAF only control wells (no cells), and positive control wells (2 µM DAF, 10 µM PAPA NONOate, in PBS). The plate was incubated at 37°C in a TECAN Infinite 200 PRO plate reader, with fluorescence intensity of each well measured each minute (using orbital averaging) for 30 min at λ_ex_ 495 nm and λ_em_ 515 nm. At the end of the assay, the wells were emptied and replaced with 100 µL of PBS, and frozen before quantification of protein content for each well using the Bradford assay. The data was analysed by calculating the linear rate of DAF fluorescence increase for each well, which was then corrected for protein content and the rate of the positive control samples. This was done to ensure reproducibility between different plates and experimental days.

### PC assay

Dinitrophenylhydrazine (10 mM in 0.5 M phosphoric acid, 100 µL per well) was added to wells containing 50 µL of cell lysate (prepared in PBS) and 50 µL of 18.2 MΩ water and incubated for 10 min at room temperature in the dark. To each well, 50 µL of 6 M sodium hydroxide was added, and the plate was incubated for 10 min at room temperature, followed by measurement of absorbance at 450 nm using a Tecan infinite 200 mx-pro plate reader. Protein carbonyl concentrations were determined for blank corrected samples using the Beer-Lambert law (molar extinction coefficient = 33131 M^-1^cm^-1^), followed by normalization to the protein content of each lysate.

### Enzymatic antioxidant activity assays

Superoxide dismutase enzyme activity was quantified using the Abcam SOD activity assay kit (Ab65354, Abcam, UK) as directed by the manufacturer. Catalase activity was determined using the Abcam catalase activity assay kit (Ab83464, Abcam, UK) as directed by the manufacturer. Enzyme activity (measured in units) was corrected for the protein concentration of samples (determined by Bradford assay as described below) for both enzyme activities.

### Bradford assay

Cell lysates (5 µL per well) were added to a 96 well plate, along with BSA standards (0-1.4 mg/mL in PBS, 5 µL per well) in triplicate. Bradford reagent (200 µL per well) was added to these samples and standards and the plate was incubated for 45 min in the dark at room temperature. Absorbance at 595 nm was measured using a Tecan infinite200 mx-pro plate reader. Protein concentration per µL of cell lysate was determined for each sample using the calibration curve of BSA standards.

### Statistical analysis

All data was checked for normality of distribution using a Shapiro-Wilks test. Data which was normal was analysed using an independent t-test or one-way ANOVA as indicated in the figure legends. Non-parametric data was analysed using either a Mann-Whitney U or Kruskal-Wallis test as indicated in the figure legends.

## Results

### Aged myotubes show morphological changes compared to young myotubes

Myotube width and length were compared for young and aged cultures (Figure 1). Aged myoblasts were significantly thinner (Figure 1A, p<0.001, unpaired t-test) and significantly shorter (Figure 1B, p<0.001, Mann-Whitney U test) than young myotubes. This confirmed that, under the culture conditions used in this study, the aged myotubes showed a dysfunctional phenotype compared to young cells.

**Figure 1:**
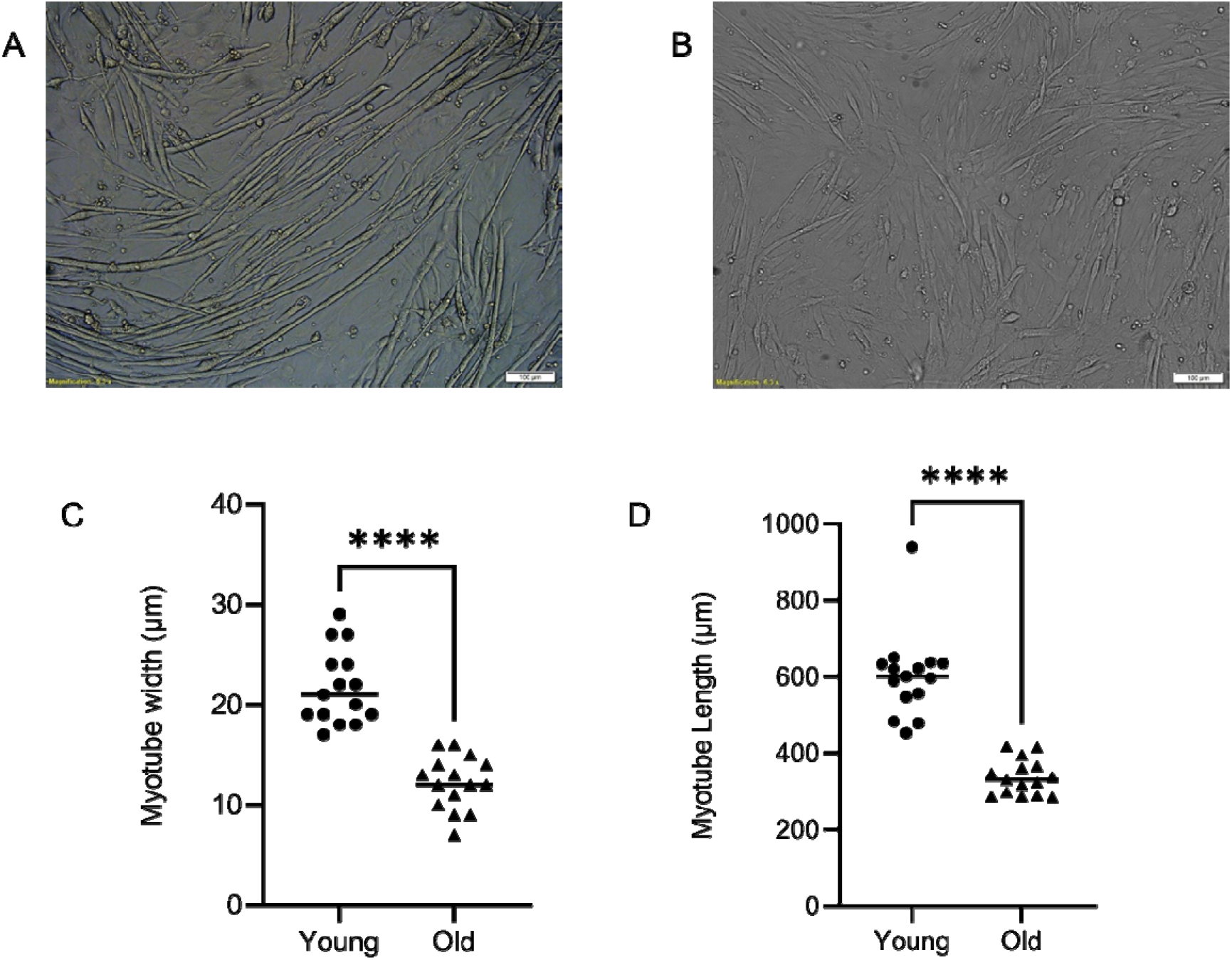
Effect of ageing protocol on myotube width and length. Representative images of young (A) and aged (B) myotubes used in analysis of myotube length and width. Scale bars equate to 100 µm. Quantification of myotube width (C) and length (D) were significantly different between young and aged. Individual replicate experiments are represented by each data-point on the graph, with the median value indicated by the horizontal line. Young and aged datasets were compared using unpaired t-test or Mann-Whitney U test, with **** p<0.001). Panel A shows myotube width (n=15 per group) and panel B shows myotube length (n=15 per group).

### Aged myoblasts show increased markers of RONS and oxidative stress compared to young myoblasts

Both young and aged myoblasts were assessed for markers of ROS production, lipid oxidation and protein oxidation (Figure 2). ROS was significantly increased in aged myoblasts compared with young myoblasts by DHE assay (Figure 2A and 1B, p<0.01 and p<0.05 respectively, unpaired t-test). Two markers of lipid oxidation were assessed (FOX assay and TBARS assay), with both markers showing a median increase in aged (compared with young) myoblasts (Figure 2C and D, p>0.05 for FOX assay, p<0.05 for TBARS assay). Protein carbonyls (measured using dinitrophenylhydrazine), as a marker of protein oxidation, showed no differences between young and aged myoblasts (Figure 2E, p>0.05, unpaired t-test). Nitric oxide levels, measured by the DAF assay, were also increased in aged myoblasts compared to young myoblast cultures (Figure 2F, p<0.05, unpaired t-test).

**Figure 2:**
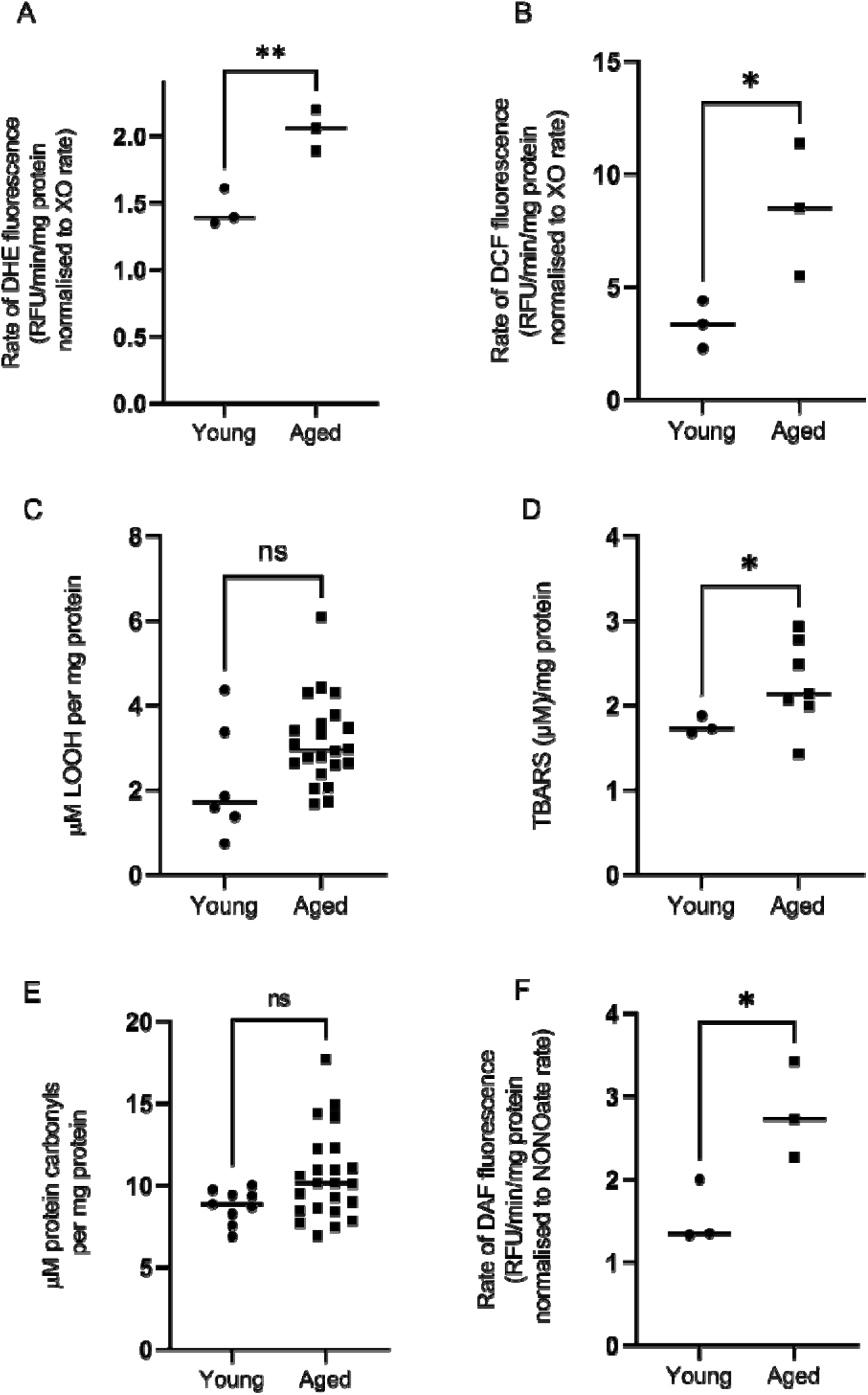
Characterisation of RONS signaling in young and aged myoblasts. Individual replicate experiments are represented by each data-point on the graph, with the median value indicated by the horizontal line. Young and aged datasets were compared using unpaired t-tests, with * p<0.05 and **p<0.01) Panel A: ROS levels were assessed in young and aged myoblast cultures by DHE fluorescence. Panel B: ROS levels were assessed in young and aged myoblast cultures by DCF fluorescence. Panel C: Lipohydroperoxides were measured by FOX assay. Panel D: Thiobuturic acid reactive substances were measured by HPLC TBARS assay. Panel E: Protein carbonyls measured using dinitrophenylhydrazine. Panel F: Nitric oxide levels measured using the DAF.

### Aged myotubes show increased markers of RONS and oxidative stress compared to young myotubes

Both young and aged myotubes were assessed for markers of ROS production, lipid oxidation and protein oxidation (Figure 3). ROS was significantly increased in aged myoblasts compared with young myotubes by DHE assay (Figure 3A, p<0.001, unpaired t-test), however this difference was not observed when ROS levels were assessed using the DCF assay (Figure 3B, p>0.05, unpaired t-test). Two markers of lipid oxidation were assessed (FOX assay and TBARS assay), with FOX assay showing a significant increase in aged compared to young, and TBARS assay showing no change in aged (compared with young) myotubes (Figure 3C and D, p>0.05 for FOX assay, p<0.05 for TBARS assay). Protein carbonyls, as a marker of protein oxidation, showed no differences between young and aged myotubes (Figure 3E, p>0.05, unpaired t-test). Nitric oxide levels were also increased in aged myotubes compared to young myotube cultures (Figure 3F, p<0.05, unpaired t-test).

**Figure 3:**
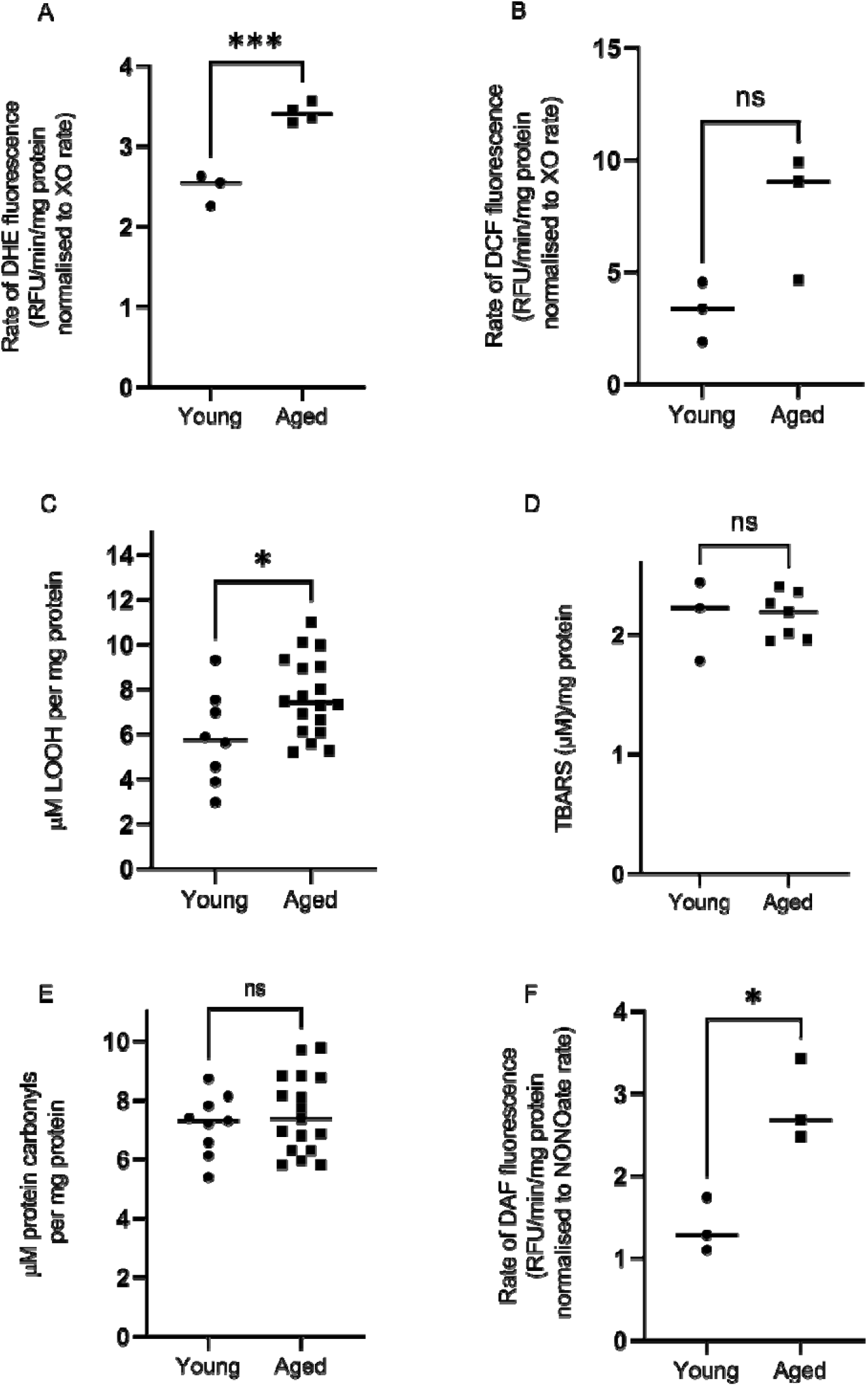
Characterisation of RONS signaling in young and aged myotubes. Individual replicate experiments are represented by each data-point on the graph, with the median value indicated by the horizontal line. Young and aged datasets were compared using unpaired t-tests, with ns p>0.05, * p<0.05 and **p<0.01) Panel A: ROS levels were assessed in young and aged myotube cultures by DHE fluorescence. Panel B: ROS levels were assessed in young and aged myotube cultures by DCF fluorescence. Panel C: Lipohydroperoxides were measured by FOX assay. Panel D: Thiobuturic acid reactive substances were measured by HPLC TBARS assay. Panel E: Protein carbonyls measured using dinitrophenylhydrazine. Panel F: Nitric oxide levels measured using DAF.

### Screening of polyphenols at physiologically-relevant exposures in aged myoblasts shows differential effects on RONS signalling

The dietary polyphenols quercetin, chrysin, curcumin, ellagic acid, kaempferol, resveratrol, epigallocatechin gallate, 3-hydroxy-tyrosol and gallic acid we assessed for effects on ROS levels in aged myoblasts (Figure 4) using the DHE assay. In aged myoblasts, curcumin and kaempferol (at 1 µM, 4 h) significantly reduced ROS production (Figure 4A, curcumin 56% ± 4%, kaempferol 54% ± 6%, p<0.05, Kruskal-Wallis test with Dunn’s multiple comparison versus solvent control). ROS production was also reduced by chrysin, ellagic acid and resveratrol, albeit not with statistical significance (Figure 4A).

**Figure 4:**
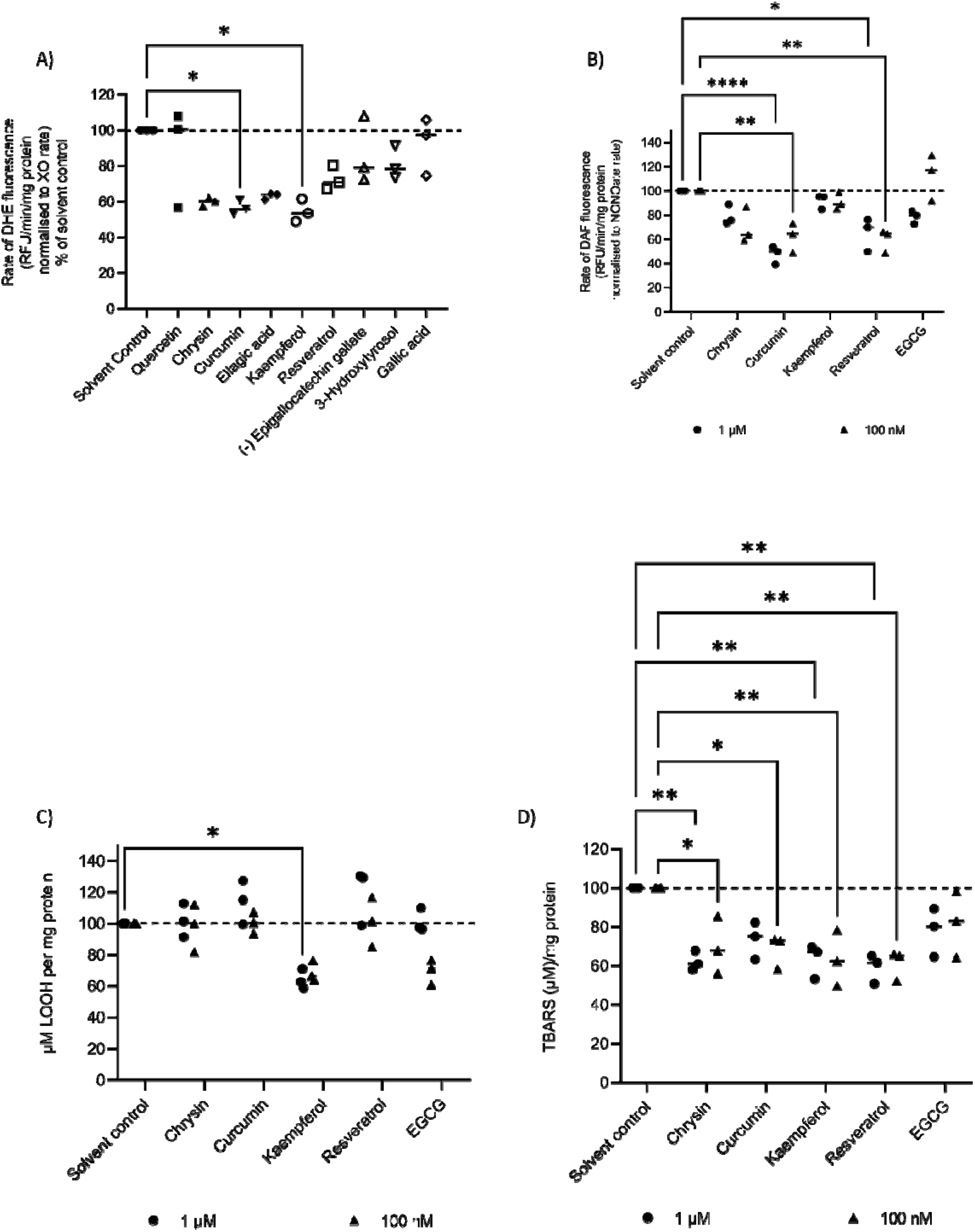
Screening of dietary polyphenols at physiologically-relevant concentrations for effects on ROS production in aged myoblasts, and characterization of hits for effects on lipid peroxidation and nitric oxide levels. Individual replicate experiments are represented by each data-point on the graph, with the median value indicated by the horizontal line. The dashed line on the graphs indicates the solvent control level. Panel A: The effect of a range of polyphenols (1 µM, 4 h) on ROS levels was determined using the DHE assay. Although several polyphenols reduced ROS levels in aged myoblasts, only curcumin and kaempferol reached statistical significance (p<0.05, Kruskal-Wallis test with Dunn’s multiple comparison versus solvent control). Panel B-D: The effects of selected polyphenols (1 µM (circles) or 100 nM (triangles), 4 h) on nitric oxide levels (DAF assay, Panel B), lipohydroperoxide concentration (Fox assay, Panel C), and TBARS (Panel D) were determined. Statistical analysis of these data used two-way ANOVA with Tukey’s multiple comparisons test.

Chrysin, curcumin, kaempferol, resveratrol and epigallocatechin gallate (EGCG) were selected for further characterisation, where the effect of exposure to these compounds (at 1 µM and 100 nM, 4 h) upon nitric oxide production (Figure 4B) and lipid peroxidation (Figure 4C and D) was determined. Nitric oxide production was significantly reduced by both curcumin (1 µM: p<0.0001; 100 nM: p<0.01) and resveratrol (1 µM: p<0.05; 100 nM: p<0.01, 2-way ANOVA with Tukey’s multiple comparisons test). Chrysin (100 nM, 4 h) also approached statistical significance (p=0.052, 2-way ANOVA with Tukey’s multiple comparisons test).

Kaempferol (at 1 µM and 100 nM, 4 h) and EGCG (100 nM, 4 h) reduced lipohydroperoxide concentrations in aged myoblasts (Figure 4C), however only kaempferol (1 µM, 4 h) reached statistical significance (p<0.05, 2-way ANOVA with Tukey’s multiple comparisons test), whereas 100 nM kaempferol and 100 nM EGCG reached near-significance (p=0.058 and p=0.068 respectively). In the TBARS assay (Figure 4D), significant reductions in lipid peroxidation were observed for chrysin (1 µM: p<0.01; 100 nM: p<0.05), curcumin (100 nM: p<0.05), kaempferol (1 µM: p<0.01; 100 nM: p<0.01), and resveratrol (1 µM: p<0.01; 100 nM: p<0.05; all the above significances tested using a 2-way ANOVA with Tukey’s multiple comparisons test).

### Screening of polyphenols at physiologically-relevant exposures in aged myotubes shows differential effects on RONS signalling and myotube morphology

The dietary polyphenols quercetin, chrysin, curcumin, ellagic acid, kaempferol, resveratrol, epigallocatechin gallate, 3-hydroxy-tyrosol and gallic acid were assessed for effects on ROS levels in aged myotubes (Figure 5) using the DHE assay. In aged myotubes, curcumin and kaempferol (at 1 µM, 4 h) significantly reduced ROS production (Figure 5A, curcumin 50% ± 9%, kaempferol 49% ± 2%, p<0.05, Kruskal-Wallis test with Dunn’s multiple comparison versus solvent control). ROS production was also reduced by chrysin, ellagic acid and resveratrol, albeit not with statistical significance (Figure 5A).

**Figure 5:**
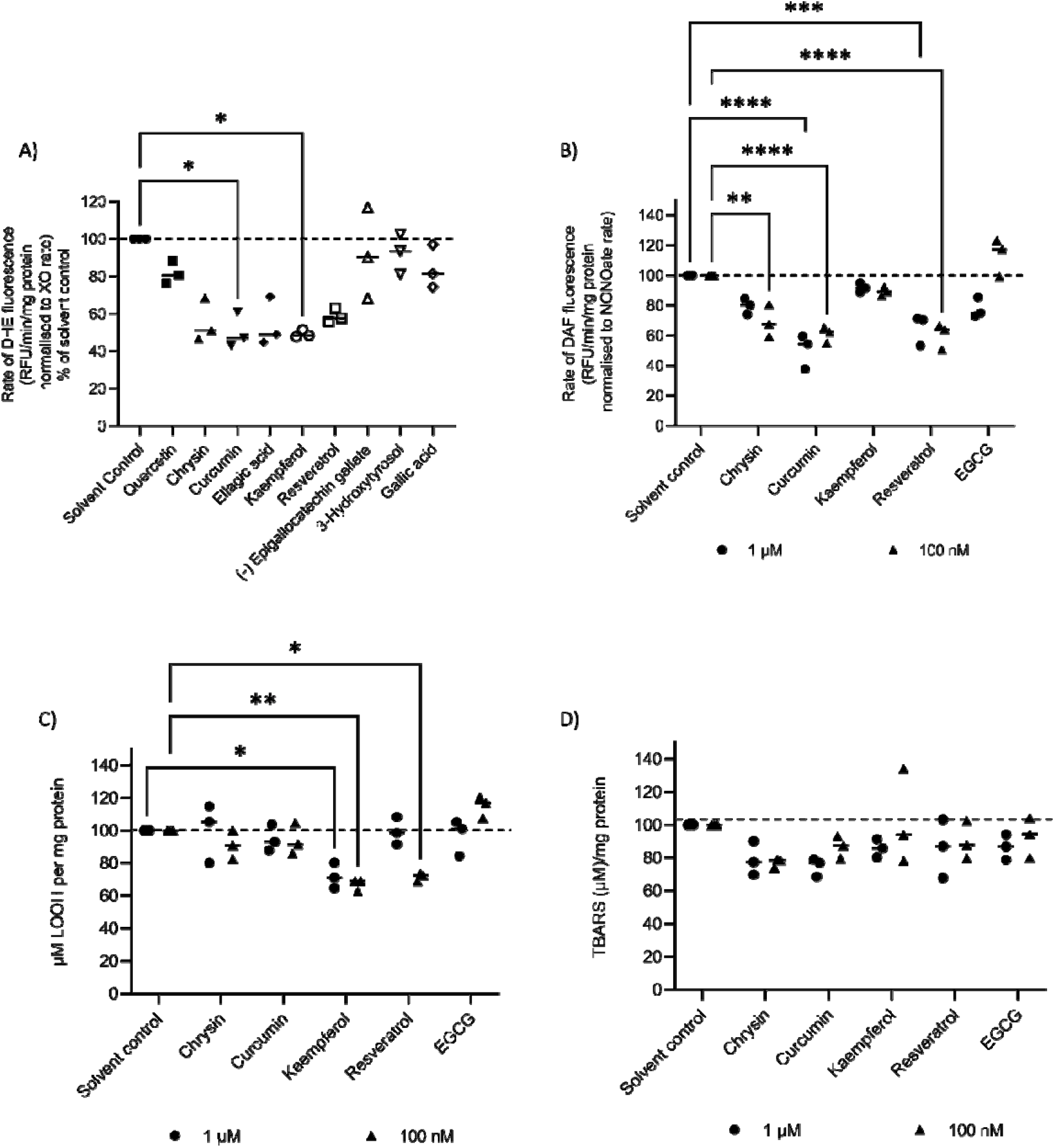
Screening of polyphenols in aged myotubes for effects on ROS levels, and characterization of hits. Individual replicate experiments are represented by each data-point on the graph, with the median value indicated by the horizontal line. The dashed line on the graphs indicates the solvent control level. Panel A: The effect of a range of polyphenols (1 µM, 4 h) on ROS levels was determined using the DHE assay. Although several polyphenols reduced ROS levels in aged myoblasts, only curcumin and kaempferol reached statistical significance (p<0.05, Kruskal-Wallis test with Dunn’s multiple comparison versus solvent control). Panel B-D: The effects of selected polyphenols (1 µM (circles) or 100 nM (triangles), 4 h) on nitric oxide levels (DAF assay, Panel B), lipohydroperoxide concentration (Fox assay, Panel C), and TBARS (Panel D) were determined. Statistical analysis of these data used two-way ANOVA with Tukey’s multiple comparisons test. Statistical significance is indicated in the main text.

Chrysin, curcumin, kaempferol, resveratrol and epigallocatechin gallate (EGCG) were selected for further characterisation, where the effect exposure to these compounds (at 1 µM and 100 nM, 4 h) upon nitric oxide production (Figure 5B) and lipid peroxidation (Figure 5C and D) was determined. Nitric oxide production was significantly reduced by chrysin (100 nM: p<0.01), curcumin (1 µM: p<0.0001; 100 nM: p<0.0001) and resveratrol (1 µM: p<0.001; 100 nM: p<0.0001, 2-way ANOVA with Tukey’s multiple comparisons test). Kaempferol (at 1 µM and 100 nM, 4 h) and resveratrol (100 nM, 4 h) reduced lipohydroperoxide concentrations in aged myotubes (Figure 5C) with statistical significance (kaempferol 1 µM: p<0.05; 100 nM: p<0.01; resveratrol 100 nM: p<0.05; 2-way ANOVA with Tukey’s multiple comparisons test). In the TBARS assay (Figure 5D), no significant reductions in lipid peroxidation were observed.

### The modulation of ROS and macromolecule oxidation in aged skeletal myotubes by dietary polyphenols is not due to upregulation of enzymatic antioxidant capacity

One potential mechanism of action for the observed effects of acute polyphenol treatment is via the upregulation of cellular antioxidant defences. To test this hypothesis, superoxide dismutase (SOD) and catalase (CAT) activities were assessed in myotube cultures (Figure 6). Statistically significant decreases in SOD (Figure 6A, p<0.01, unpaired t-test) and CAT (Figure 6B, p<0.05, unpaired t-test) activities were observed when comparing young and aged myotubes, suggesting that cellular antioxidant defence may be compromised in aged myotubes. Dietary polyphenol treatment of aged myotubes did not alter either SOD (Figure 6C) or CAT (Figure 6D) activities, suggesting upregulation of cellular enzymatic antioxidant capacity is not part of the mechanism of action for the observed reduction of ROS and lipid oxidation in aged myotubes.

**Figure 6:**
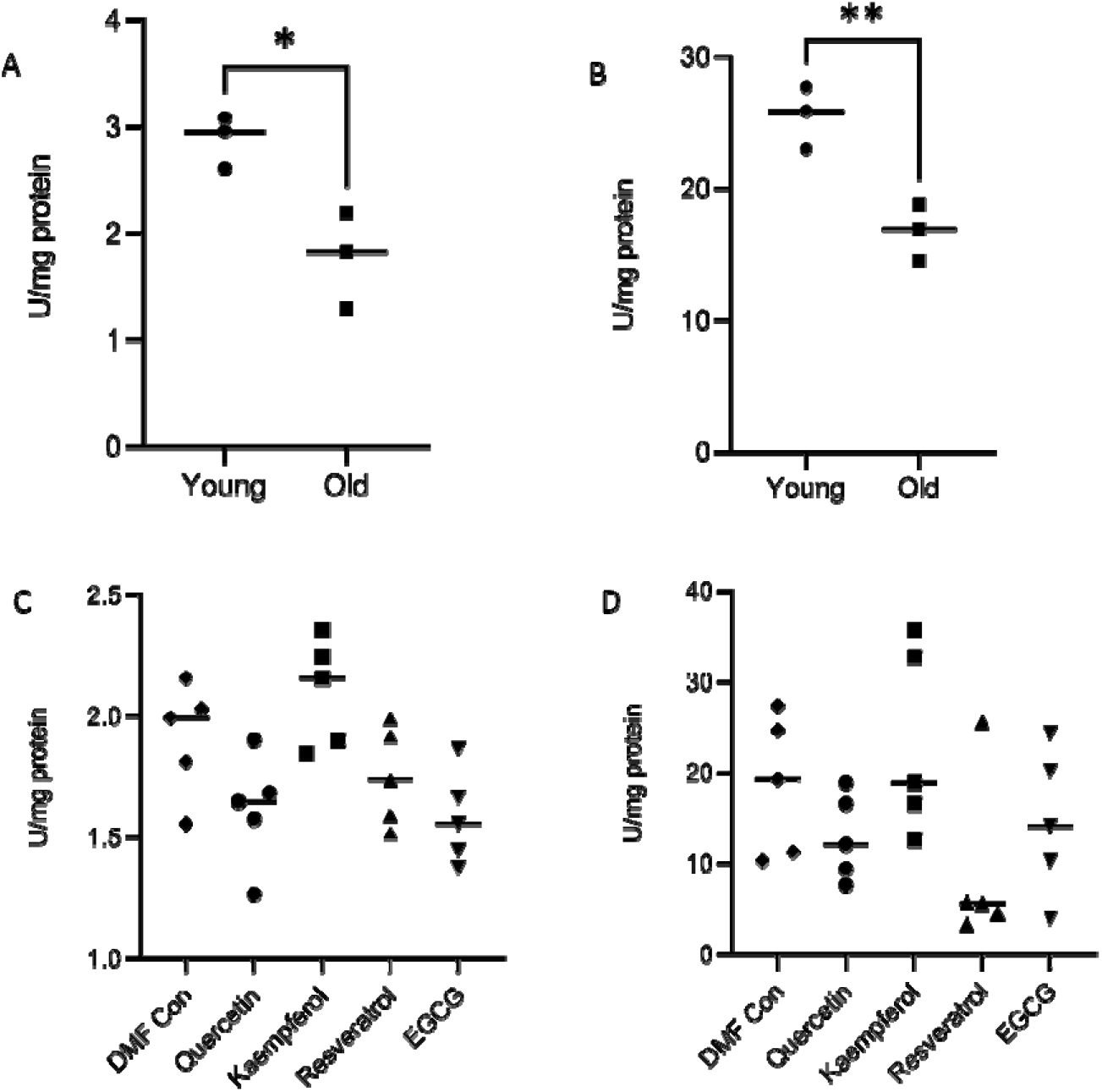
Superoxide dismutase and catalase activities in aged myotubes are not modulated by polyphenol exposure. Panel A and C show catalase activity assays, panel B and D show superoxide dismutase (SOD) assays. Both catalase (A) and SOD (B) activities are significantly reduced in aged myotubes compared to young myotubes (* p<0.05, **p<0.01, unpaired t-test). Polyphenol exposure (100 nM, 4 h) does not significantly alter Catalase (C) or SOD (D) activities in aged myotubes (p>0.05, one-way ANOVA and Kruskal-Wallis test respectively). Individual replicate experiments are represented by each data-point on the graph, with the median value indicated by the horizontal line.

### Acute kaempferol exposure of aged myoblasts partially induces a young phenotype when differentiated into myotubes

To understand if acute exposure to these polyphenols impacted on myotube morphology, myoblasts were treated with 100 nM for 4 h of Quercetin, Kaempferol, EGCG or Resveratrol prior to differentiation into myotubes (Figure 7). Kaempferol exposure resulted in a significant (p<0.05) increase in myotube length (Figure 7C), but no change in myotube width (Figure 7D). When aged myotubes were incubated with these polyphenols after differentiation, no changes in in myotube length or width were observed (Figure 8C and 8D).

**Figure 7:**
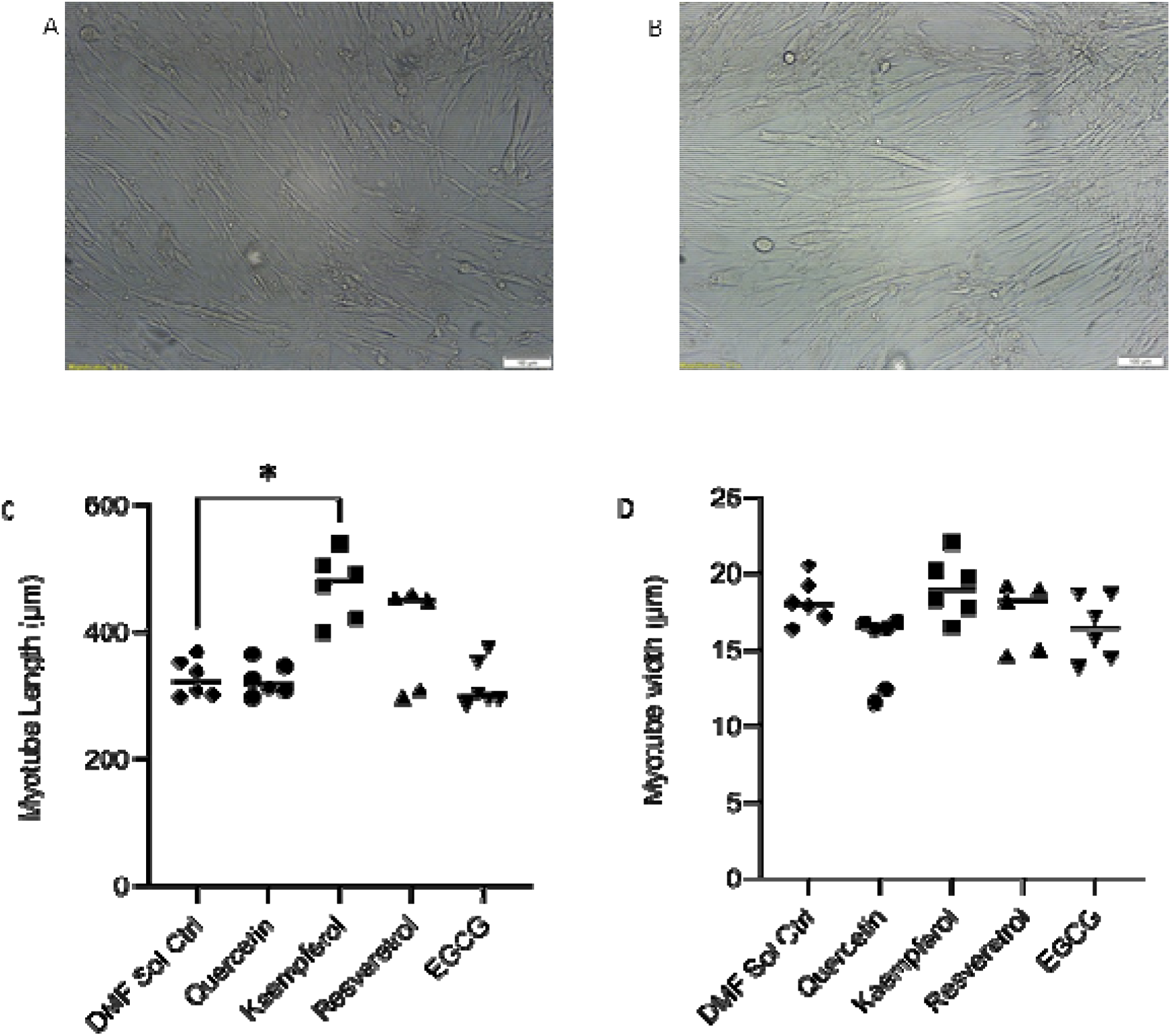
The effect of acute polyphenol exposure on myotube length and width. Panels A and B show representative images of myotubes resulting from the differentiation of solvent control (A) and kaempferol-treated (100 nM, 4 h, Panels B) aged myoblasts treated before differentiation. Scale bars indicate 100 µm. Panels C (myotube length) and D (myotube width) for solvent only or polyphenol-treated (100 nM, 4 h) myoblasts prior to differentiation. Individual replicate experiments are represented by each data-point on the graph, with the median value indicated by the horizontal line. Datasets were compared using a Kruskal Wallis test (p<0.01), with Dunn’s multiple comparisons test versus the solvent control; * p<0.05.

**Figure 8:**
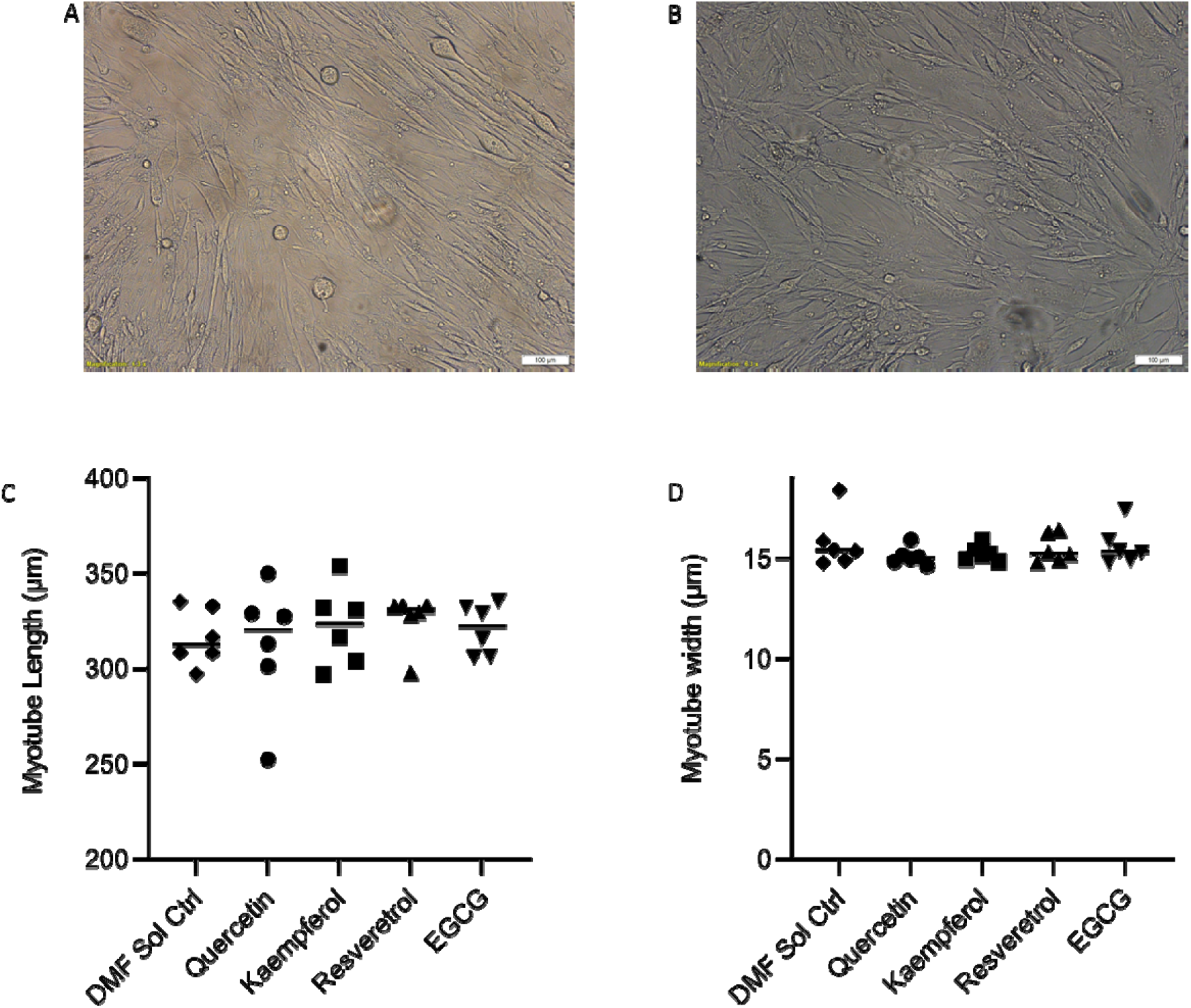
The effect of acute polyphenol exposure post differentation on myotube length and width. Panels A and B show representative images of aged myotubes treated with solvent control (A) and kaempferol-treated (100 nM, 4 h, Panels B) after differentiation. Panels C (myotube length) and D (myotube width) for solvent only or polyphenol-treated (100 nM, 4 h) myoblasts post differentiation. Individual replicate experiments are represented by each data-point on the graph, with the median value indicated by the horizontal line. Datasets were compared using a Kruskal Wallis test (p>0.05).

## Discussion

Although oxidative stress endpoints have been previously assessed in C_2_C_12_ cells, this study is the first (to authors knowledge) to undertake a thorough assessment of RONS signaling changes between young and aged (by multiple population doublings) myoblast and myotube cell cultures, and use this model to screen for potential modifiers of RONS signaling dysfunction in aged skeletal muscle. Previous studies of oxidative stress in C_2_C_12_ have used a range of stressors such as TNF-_α_ (J Pharm Biomed Anal, 2021, 204, 114271), LPS (Poult Sci. 2019 Jul 1;98(7):2756-2764), and hydrogen peroxide (J Biosci Bioeng. 2021 May;131(5):572-578; Molecules 2021 Jan 4;26(1):215; J Pharmacol Exp Ther. 2021 Mar;376(3):385-396; Front Cell Dev Biol. 2020 Sep 11;8:541260), whereas this study uses repeated population doublings to mimic the ageing process. The C_2_C_12_ model used in this study has been well-characterized in a physiological context (such as alterations to myotube structure, time to differentiation from myoblasts to myotubes, maintenance of a skeletal muscle phenotype, e.g., J. Cell. Physiol., 2010, 225, 240) to reflect an aged skeletal muscle phenotype, an observation our study agrees with (based on a reduction of myotube length and width compared with young cells, Figure 1), and this study has now demonstrated that it also reflects biochemical alterations in RONS signaling, such as those reported from aged muscle biopsies. Increases in ROS levels and markers of lipid oxidation have been shown for both myoblasts and myotubes in aged cultures compared with young cultures, whereas protein carbonyl levels were not different in young and aged cultures (Figures 2 and 3).

These observations show good agreement with human biopsy data for studies comparing young and aged volunteers, showing consistent increases in lipid oxidation markers (biopsy data is quite variable for protein carbonyl level, with some studies showing differences and others no changes) supporting the validity of the myoblast and myotube culture models (Free Rad. Biol. Med., 1999, 26, 303; Redox Biol., 2015, 5, 267; Free Rad. Biol. Med., 2006, 41, 797). Increases in nitric oxide levels have also been reported in aged skeletal muscle samples from rodent models (e.g., Free Radic Biol Med. 2015 Jan;78:82-8) and 3-nitrotyrosine (a marker of protein nitrosylation and surrogate marker for nitric oxide) has been reported to increase in human biopsy samples from aged compared with young participants (Cells 2019, 8(12), 1525). Taken together, these data suggest that the C_2_C_12_ model used in this study is representative of aged skeletal muscle biopsy samples in both phenotypic and RONS signalling contexts.

A major limitation of skeletal muscle ageing and sarcopenia research is a lack of a tractable model to screen compounds for potential beneficial effects on muscle ageing and dysfunction. Although there are a range of *in vivo* model systems (e.g., murine and other rodent models), these lack the throughput and capacity to screen a large number of compounds. This study demonstrates that the C_2_C_12_ ageing model can address this issue. Both aged myoblast and myotube cultures were used as a screening tool to identify dietary polyphenols with bioactivity on RONS signaling from a wider panel of polyphenols. Chrysin, curcumin, kaempferol and resveratrol were identified to reduce elevated markers of RONS levels and lipid peroxidation at physiologically-relevant exposures. Kaempferol also had a significant effect on myotube length, when incubated with aged myoblasts prior to differentiation (Figure 7), which is representative of a younger phenotype in this cell model of ageing. This screening approach, combined with the use of compound exposure concentrations and durations guided by known pharmacokinetic parameters for these compounds, are two areas in which this work has advanced the field.

Resveratrol, curcumin, epigallocatechin gallate and quercetin have all been previously assessed for effects on skeletal muscle aging in cell culture and rodent models (Int J Mol Sci, 2019, 20, 1178; J Agric Food Chem, 2021, 69, 6214; FASEB, 2014, 28, 1159.4; Antioxidants, 2021, 10, 476; Protein and Cell, 2019, 10, 770; Braz. Pharm Sci, 2020, 56; J Frail Ageing, 2015, 4 209; Exp Gerontol, 2008, 43, 176; Arch Biochem Biophys, 2020, 692, 108511; Biochem and Biophys Res Comm, 2017, 489, 142). These studies have shown that these polyphenols alter ROS signaling in skeletal muscle cells and lysate, and some, but not all, rodent studies report improvements in muscle function with chronic exposures. A major limitation of these studies is that the *in vitro* experiments use exposures which do not reflect human pharmacokinetic data (for example, Chang et al exposed C_2_C_12_ cells to 50 µM epigallocatechin gallate for 24 h, whereas the reported Cmax and half-life for epigallocatechin galate in humans is 447 ± 300 nM and 3.6 ± 1.4 h (Arch Biochem Biophys, 2020, 692, 108511, Cancer Epidemiol Biomarkers Prev. 2002, 1025-32.). Additionally, even the rodent model studies highlighted above use impractical doses of polyphenols when scaled to human studies (for example, both Braz. Pharm Sci, 2020, 56, and J Frail Ageing, 2015, 4 209-215 dosed rats with 200 mg/kg body weight per day, which when scaled to a 75 kg human (as detailed in J Basic Clin Pharm, 2016, 7, 27-31, taking body surface area into consideration) equates to a bolus of 2.43 g of pure quercetin or epigallocatechin gallate per day respectively. Human clinical trials using quercetin report minor side effects at long term doses over 1g, (J Am Heart Assoc, 2016, 5, e002713), suggesting that the exposures used in the rodent models are not reflective of what would be used in humans. This lack of relevance to human pharmacokinetics in both in vitro and animal model experiments means that it is difficult to extrapolate these findings to humans. This study has, for in vitro models, used more pharmacokinetically-relevant exposures, using a 4 h exposure at 1 µM for initial screening, and a lower concentration (100 nM, 4 h) which is also consistent with reported pharmacokinetic data for the test compounds (Cancer Epidemiol Biomarkers Prev. 2008 Jun;17(6):1411-7; Cancer Epidemiol Biomarkers Prev. 2002, 1025-32; Br. J. Clin. Pharm., 51 (2) (2001), pp. 143-146; Ann N Y Acad Sci. 2011 Jan;1215:9-15; Nutrients 2019, 11(10), 2288).

This study has also undertaken some limited mechanistic investigations, to try to understand how dietary polyphenols such kaempferol modulate RONS signalling, via the assessment of enzymatic antioxidant activities (SOD and catalase) in aged myotube cultures (Figure 6). Kaempferol has been reported in a variety of studies to modulate SOD and catalase activity in a variety of health and disease states (Drug and chemical toxicology, 39(3), 239–247; Oxidative Medicine and Cellular Longevity, vol. 2018, Article ID 1610751; Redox Report, 20:5, 198-209). It should be noted, however, that the application of physiologically-relevant kaempferol exposures in these studies is variable. This study reports that aged myotubes have reduced SOD and catalase activities compared to young myotubes, and that none of the polyphenols tested induced either SOD or catalase activities in aged myotube cultures. This suggests that modulation of enzymatic antioxidant capacity is not the mechanism of action of the polyphenols tested which underpins the reduction in ROS and lipid peroxidation markers reported above (Figures 4 and 5).

It should be noted that there are some potentially important limitations with the experimental design in this study. Dietary polyphenols undergo significant conjugative metabolism (e.g., glucuronidation, sulfation, methylation), however due to a lack of availability of human polyphenol metabolites, they were not used in the experiments described in this study. Additionally, the experiments described above assess the effects of acute exposures to dietary polyphenols, whereas a chronic dosing is more likely to be experienced by humans. It should also be acknowledged that this study only assesses the direct impact of polyphenol exposure on skeletal muscle cells, and does not integrate effects on other cell types or systems (e.g., neuronal cells, blood supply).

## Conclusions

Overall, this study has characterized a myoblast and myotube cell culture model of skeletal muscle ageing for changes in RONS signaling. These cultures show similar changes to those reported for human skeletal muscle biopsies, and the modulation of RONS signaling by physiologically-relevant dietary polyphenol exposures in aged myoblasts and myotubes has been demonstrated. This model system can be used to identify bioactive compounds which may be beneficial for restoring normal RONS signaling in aged skeletal muscle.

## Acknowledgements

NH was funded by a University of Hull PhD studentship. We thank Dr Roger Sturmey and Mr Andrew Gordon for their assistance with the TBARS assay. A preprint of this work has previously been published at bioRxiv (bioRxiv 2021.11.20.469396)

## Author statements

All the authors declare that there are no conflicts of interest with this research.

HSJ: Designed the study, planned experiments, analysed data, wrote the manuscript.

NH: Designed the study, planned experiments, analysed data, wrote the manuscript.

MF: Designed the study, wrote the manuscript

LS: Designed the study, wrote the manuscript.

